# Building and analyzing metacells in single-cell genomics data

**DOI:** 10.1101/2024.02.04.578815

**Authors:** Mariia Bilous, Léonard Hérault, Aurélie AG Gabriel, Matei Teleman, David Gfeller

## Abstract

The advent of high-throughput single-cell genomics technologies has fundamentally transformed biological sciences. Currently, millions of cells from complex biological tissues can be phenotypically profiled across multiple modalities. The scaling of computational methods to analyze such data is a constant challenge and tools need to be regularly updated, if not redesigned, to cope with ever-growing numbers of cells. Over the last few years, metacells have been introduced to reduce the size and complexity of single-cell genomics data while preserving biologically relevant information. Here, we review recent studies that capitalize on the concept of metacells – and the many variants in nomenclature that have been used. We further outline how and when metacells should (or should not) be used to study single-cell genomics data and what should be considered when analyzing such data at the metacell level. To facilitate the exploration of metacells, we provide a comprehensive tutorial on construction and analysis of metacells from single-cell RNA-seq data (https://github.com/GfellerLab/MetacellAnalysisTutorial) as well as a fully integrated pipeline to rapidly build, visualize and evaluate metacells with different methods (https://github.com/GfellerLab/MetacellAnalysisToolkit).

## Introduction

As single-cell sequencing becomes more affordable and widely used, both the number and the size of single-cell genomics datasets are growing exponentially^1,2^ with no signs of slowing down. Current single-cell sequencing studies from tissues, organs and whole organisms can profile millions of cells^3–9^. Along with the development of integration algorithms, even larger atlases have been built, reaching tens of millions of cells^10^. Historically, most of the single-cell genomics studies have focused on transcriptomic profiling (scRNA-seq). More recently, to expand our understanding of cellular complexity, other modalities have been integrated in single-cell genomics technologies, including epigenomics (e.g., scATAC-seq)^11^, surface proteins (e.g., CITE-seq)^12^, and adaptive immune receptor (AIR) in B cells (i.e., BCR sequences) or T cells (i.e., TCR sequences)^13,14^. Moreover, simultaneous profiling of multiple modalities at single-cell resolution can be performed nowadays to discover novel cell types^15^, better characterize known cell types^16^, identify gene regulatory interactions^17^ and biomarkers^16,18^, and recover comprehensive immune repertoire^19^.

The majority of single-cell genomics datasets are generated using droplet-based sequencing technologies^2,20–23^. These techniques have very high throughput, allowing researchers to profile millions of cells, but relatively low sensitivity due to low-depth sequencing and limited efficiency of the retro-transcription/amplification procedure. As a result, many transcripts are missed, something referred to as dropout or non-biological zeros in the single-cell profile matrices.

Simultaneous profiling of millions of cells across multiple modalities provides unprecedented opportunities to map the full heterogeneity of whole organs in health and diseases but generates computational challenges to analyze such large-scale data. Several approaches have been designed to cope with the size and inherent noise of single-cell data. Both hardware and software developments enable users to analyze very large datasets, although this often comes with a price in terms of speed and practicality (e.g., the need to use dedicated machines with very high memory). Different downsampling and sketching scenarios have been designed to reduce the size of the data^24–26^. In parallel, multiple approaches have been introduced to address the dropout issue, including imputation^27–33^ or customizing algorithms to zero-inflated data^34–37^. More recently, metacells – defined as groups of highly similar cells which are aggregated together – have been proposed as a way to simultaneously reduce the size and improve the signal-to-noise ratio in single-cell genomics data^38–41^.

Here, we review studies that have introduced the metacell concept, have developed metacell construction tools and quality metrics, and have used metacells for single-cell sequencing data analysis. We discuss the pros and cons of using metacells and provide recommendations for building metacells and analyzing single-cell data at the metacell level. These recommendations are accompanied by a comprehensive tutorial (https://github.com/GfellerLab/MetacellAnalysisTutorial) as well as an integrated pipeline allowing users to build and analyze metacells with different tools (https://github.com/GfellerLab/MetacellAnalysisToolkit).

## Metacell concept

Metacells are defined as a partition of single-cell data into disjoint homogeneous groups of highly similar cells followed by aggregation of their profiles (Fig.1). This concept relies on the assumption that most of the variability within metacells corresponds to technical noise and not to biologically relevant heterogeneity. As such, metacells aim at removing some of the noise while preserving the biological information of the single-cell data. The metacell concept was introduced in 2019 by Baran and colleagues in ref.^38^ with the motivation to obtain robust profiles in sparse scRNA-seq data. Roughly at the same time, Iacono and colleagues in ref.^42^ proposed to aggregate similar cells into metacells (named ‘iCells’ in the original study) as a way to overcome the computational burden associated with the analysis of large scRNA-seq data within the bigSCale computational framework^42^. These two pioneering studies reflect two main aspects of the usage of metacells: (i) enhancing signal in sparse scRNA-seq data and (ii) lowering computational burden due to the large size of single-cell genomics data. Since then, several other studies have built upon the metacell concept^39–41^, extending its application to other single-cell modalities, including scATAC-seq^41^, CyTOF^43^, FACS, AIR^44^, as well as multimodal single-cell data^45^.

**Figure 1.**
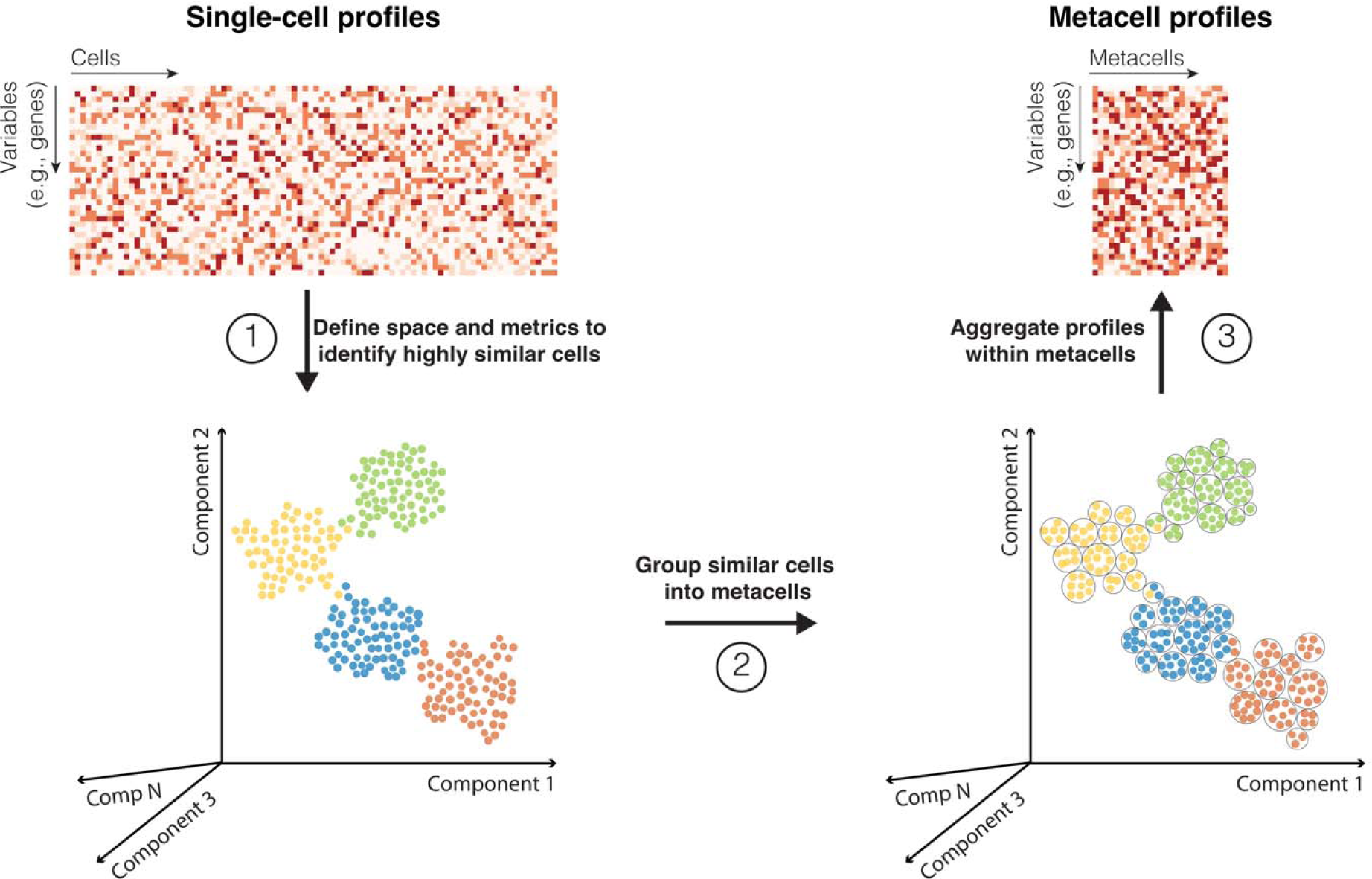
Main conceptual steps in the metacell construction workflow. Starting from a single-cell profile matrix, space and metrics are first defined for identifying cells displaying high similarity in their profiles (e.g., high transcriptomic similarity in scRNA-seq data). Second, highly similar cells are grouped into metacells. Third, single-cell profiles within each metacell are aggregated to create a metacell profile matrix. Dots represent single cells colored by cell type.

### Building metacells

Different tools have been developed and are currently available to build metacells (see Box 1). The first version of the MetaCell algorithm by Baran and colleagues used multiple resampling of the k-nearest neighbor (kNN) graph to identify robust groups of highly similar cells^38^. This approach was poorly scalable for large datasets and a new version of the original MetaCell tool (referred to as MC2^39^) was introduced based on a divide-and-conquer approach. In parallel, an alternative method of metacell construction and analysis, named SuperCell^40^, was developed based on a graph-based clustering approach. Later on, a metacell tool named SEACells^41^, was proposed based on the concept of archetypes^46^. These methods follow the original idea of metacells and use the same nomenclature. Technical details on how groups of highly similar cells are identified and aggregated in these different metacell construction tools are available in Box 1.

##### Box 1. Overview of several tools providing standalone metacell construction pipelines.

**MetaCell (implemented in R).** A single-cell kNN graph is built from the correlation of log-normalized counts based on a set of highly variable genes. Edges of the graph are further re-weighted followed by graph re-balancing (in terms of in- and outcoming edges). The balanced graph is resampled multiple times and clustered with a graph adaptation of k-means. Then, it is re-weighted based on resampling co-occurrence results. The final metacell partition along with a list of outliers is obtained via a graph adaptation of k-means. Expression profiles of metacells are computed as a mean of counts within metacells normalized by a median averaged gene count across metacells followed by logarithmic scaling. The downstream analysis is performed with a MetaCell framework and the metacell object can be converted to a single-cell-experiment (SCE)^59^ object.

**MetaCell2 (MC2, implemented in Python).** To accelerate the original MetaCell algorithm, MC2 applies a two-phase divide-and-conquer approach. Firstly, cells are randomly divided into piles of ∼10k cells and initial metacells are built applying a MetaCell^38^-like approach per pile. Then, transcriptionally similar metacells are grouped into metagroup piles for the identification of final metacells and outliers. The profiles of metacells are computed by summing up counts within metacells followed by normalization by the total number of counts in a metacell. The downstream analysis is performed with Scanpy^60^ or Seurat^26,61^ frameworks. To recover very rare cell types, MC2 has two approaches. In the first approach, outlier cells are allocated into a separate pile and metacells are re-computed under the assumption that rare cells could be scattered among different piles and could not form their own metacell. In the second approach, very rare gene modules are identified from the entire dataset. Then, a separate pile of cells with high expression of rare gene modules is generated and analyzed for metacell partition.

**SuperCell (implemented in R).** A single-cell kNN graph is built from the Euclidean distances between cells in the principal component space. Metacells are computed by applying the walktrap community detection algorithm^62^. Given its hierarchical structure, this approach allows users to probe at once all possible numbers of metacells (i.e., graining levels). By default, the normalized counts are averaged within metacells. Alternatively, raw counts can be summed. The downstream analysis can be performed using the SuperCell framework, which accounts for metacell size (i.e., number of single cells in a metacell), or using conventional Seurat^26,61^ or SCE^59^ pipelines (not accounting for metacell size). To scale the method for larger datasets, an approximate solution has been proposed where metacells are identified on a subset of cells and the rest of the cells are mapped to the closest metacells.

**SEACells (implemented in Python).** A single-cell kNN graph is built from the Euclidean distance in the principal component (or singular value decomposition for scATAC-seq) space. Distances in the graph are transformed to affinity by applying an adaptive Gaussian kernel. The affinity matrix is then decomposed into archetype and membership matrices. The archetype matrix represents archetypes as a linear combination of cells and the membership matrix represents cells as a linear combination of archetypes. To fit decomposition, the archetype matrix is initialized with waypoints using a maximum minimum sampling approach^63^, ensuring more representative initialization. Single cells are assigned to a given metacell based on the maximum membership value of the corresponding archetype. Raw counts are summed up. The downstream analysis is performed with a standard Scanpy framework^60^.

Other tools use metacells as part of a broader pipeline or method designed for different purposes (see Supplementary Table 1). Some of them, including iCells^42^, msPHATE^44^, scCorr^47^, scWGCNA^48^, popInfer^49^, and C ^50^, developed a dedicated method for metacell identification. Others use existing tools or compute metacells by excessive clustering of the data^43,51–56^. Of special interest is the multiscale PHATE (msPHATE)^44^ approach for data representation at multiple resolutions, which can be considered as a tool for metacell construction, whereby highly similar cells are aggregated into metacells (named ‘groups’ in the original study) using the approach of diffusion condensation^57^. This study was among the first to demonstrate that the concept of metacells, initially proposed for scRNA-seq data, can be applied to scATAC-seq, flow cytometry, CyTOF, AIR and even clinical data. Many of these studies utilize alternative nomenclature for metacells, including iCells^42^, supercells^43^, clusters^47^, pseudocells^48,49,52,55^, miniclusters^51^, micro-clusters^53^, micro-states^54^, groups^44^ or cellstates^50^. To avoid ambiguities, we encourage future studies to use the well-established term ‘metacell’ originally coined in Baran et al.^38^.

Most of the existing methods for metacell construction rely on single-cell similarity/affinity graph partitioning^38–41,44^ and further aggregation of nodes grouped together. This makes metacell construction related to the concept of graph coarse-graining^57,58^.

Once metacells are identified, their profiles are computed by aggregating the profiles of the single cells belonging to each metacell. The aggregation is usually done by either summing raw counts^38,42,41^ or by averaging normalized counts^40,44,47^. The first approach is recommended if metacells are to be used for comprehensive downstream analyses, while the second approach is more suitable when metacells are used for some specific analyses (see ‘Workflow recommendations’ section).

### Graining level

A key element in metacell construction is the choice of the number of metacells. The ratio between the number of single cells and the number of metacells determines the level of size reduction between the single-cell and the metacell data. This ratio is referred to as the graining level^40^. A graining level of 1 corresponds to a single-cell resolution and a graining level of 50 corresponds to a 50-fold reduction in the number of metacells compared to single cells in the initial dataset (Fig. 2a, see Materials and Methods for the description of the datasets used throughout this review). Given that metacells are defined as a disjoint partition of data, the graining level corresponds to the average size of metacells (i.e., the average number of single cells in metacells).

**Figure. 2.**
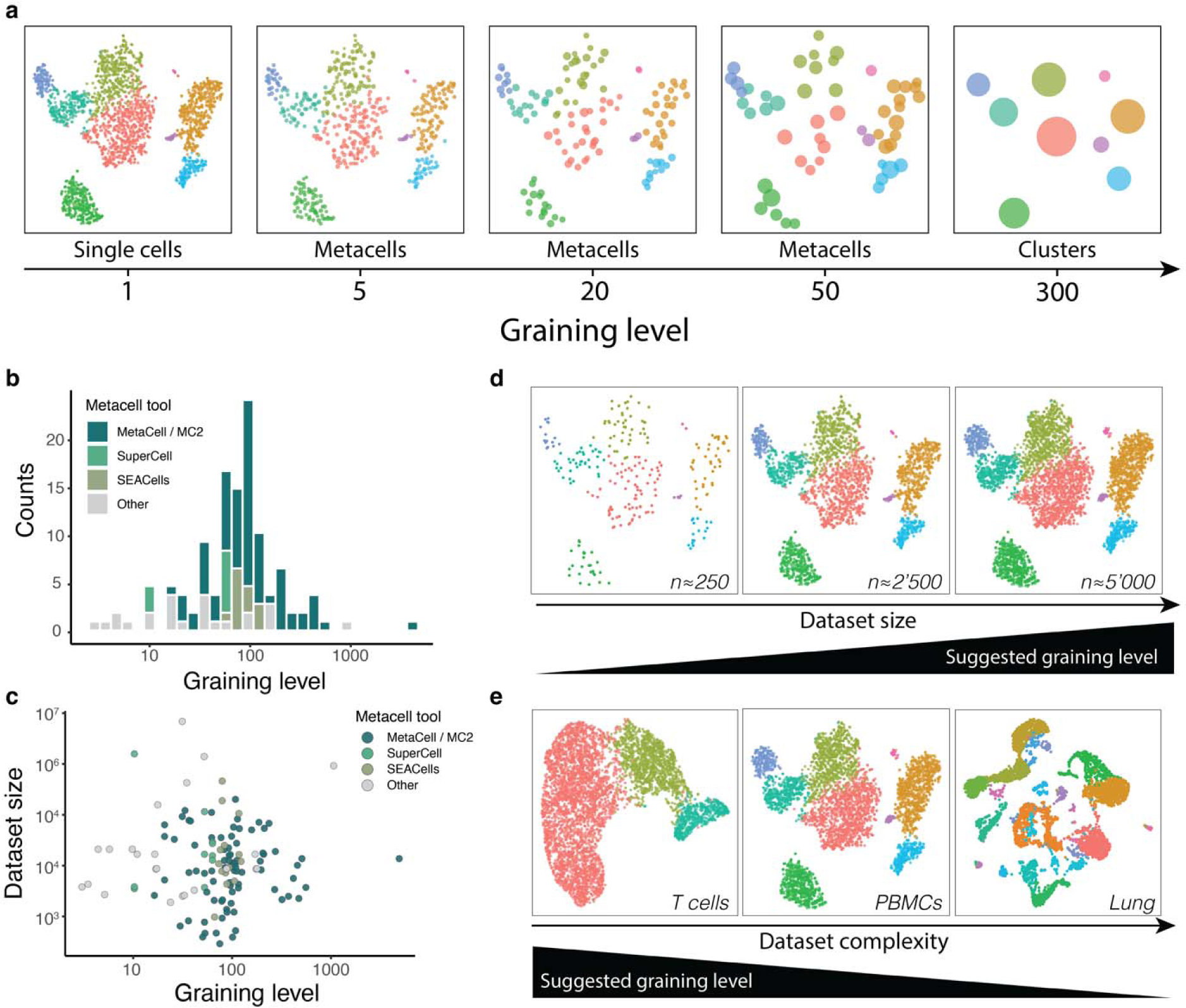
Graining level of metacell partition. **a**, tSNE^64^ representation of a PBMC scRNA-seq dataset (see Materials and Methods) at different graining levels. Each dot represents a single cell, a metacell or a cluster, depending on the graining level. Colors represent cell types. **b**, Distribution of graining levels in different studies using metacells (see Supplementary Table 2). Colors represent different metacell construction tools. **c** Graining levels used for datasets of different sizes. Colors represent different metacell construction tools. **d-e**, The choice of a graining level is influenced by the size (d) and the complexity (e) of a particular dataset: larger graining levels can be applied to larger and/or less complex datasets. Colors represent cell types.

Metacell construction tools have used different approaches to address the choice of graining levels. In MetaCell and the original version of MC2 (v 0.8.0), the user specifies a minimum or targeted metacell size. Accounting for this constraint, a resulting metacell partition is obtained by iteratively optimizing internal metrics that evaluate both the stability of the resulting partition in terms of balanced connectivity and the homogeneity of metacell sizes. In SuperCell, SEACells and the latest version of MC2 (v 0.9.4), the user specifies the graining level, or equivalently the number of metacells. As such, all these metacell construction methods require users to specify either the actual graining level or parameters that will implicitly determine the graining level.

There is currently no consensus on the definition of an optimal graining level and its estimation. Some approaches suggest that there is a range of acceptable graining levels that effectively represent the initial data by preserving the overall and cell-state-specific heterogeneity^40,41^. To get an overview of the range of graining levels that have been used, we performed a meta-analysis of several studies using metacells (Fig. 2b, Supplementary Table 2). Graining levels ranged from very small (∼5) to very large (>1000), with a median of around 80. The MetaCell algorithm resulted in larger graining levels (median at 90) compared to other methods (median at 55). There was no correlation between dataset size and the graining level used for building metacells (Fig. 2c), suggesting that an appropriate graining level also depends on dataset complexity.

When selecting a graining level for a particular dataset, we suggest considering two parameters: the dataset size and complexity. Larger gaining levels are often acceptable for larger datasets with higher redundancy (e.g., each cell type/state is covered by hundreds of single cells) (Fig. 2d). For a given size, lower graining levels should be used for datasets of higher complexity (e.g., whole organ or whole organism sequencing) as compared to datasets of lower complexity (e.g., a pre-defined cell type) to retain the underlying heterogeneity (Fig. 2e). Practically, it is recommended to choose graining levels such that the resulting number of metacells is at least ten times larger than the expected number of cell subtypes to ensure statistical power in downstream analyses at the metacell level and lower the risk of having cells of different cell subtypes aggregated in the same metacell. In most cases, a graining level somewhere between 20 and 75 satisfies these requirements, and our recent work indicates that many downstream analyses give consistent results across this range of graining levels^40^.

A graining level can be considered as reasonable if it preserves all the cell states present in the single-cell data and the biological heterogeneity within them while reducing technical noise. We also note that an ‘optimal’ graining level – if it exists – is difficult to infer with scores developed for the estimation of an optimal number of clusters (e.g., modularity^65^, silhouette coefficient^66^ or other approaches developed specifically for scRNA-seq data^67–70^) due to the differing objectives of clustering and metacells.

### Metacell quality metrics

An ideal metacell should contain highly similar cells so that the heterogeneity within a metacell is only due to technical aspects (e.g., transcript sampling error) and no biologically relevant information is lost between the single-cell and the metacell level. To quantify the quality of a metacell partition, several approaches have been developed (see Table 1 for analytical formulas).

**Table 1.**
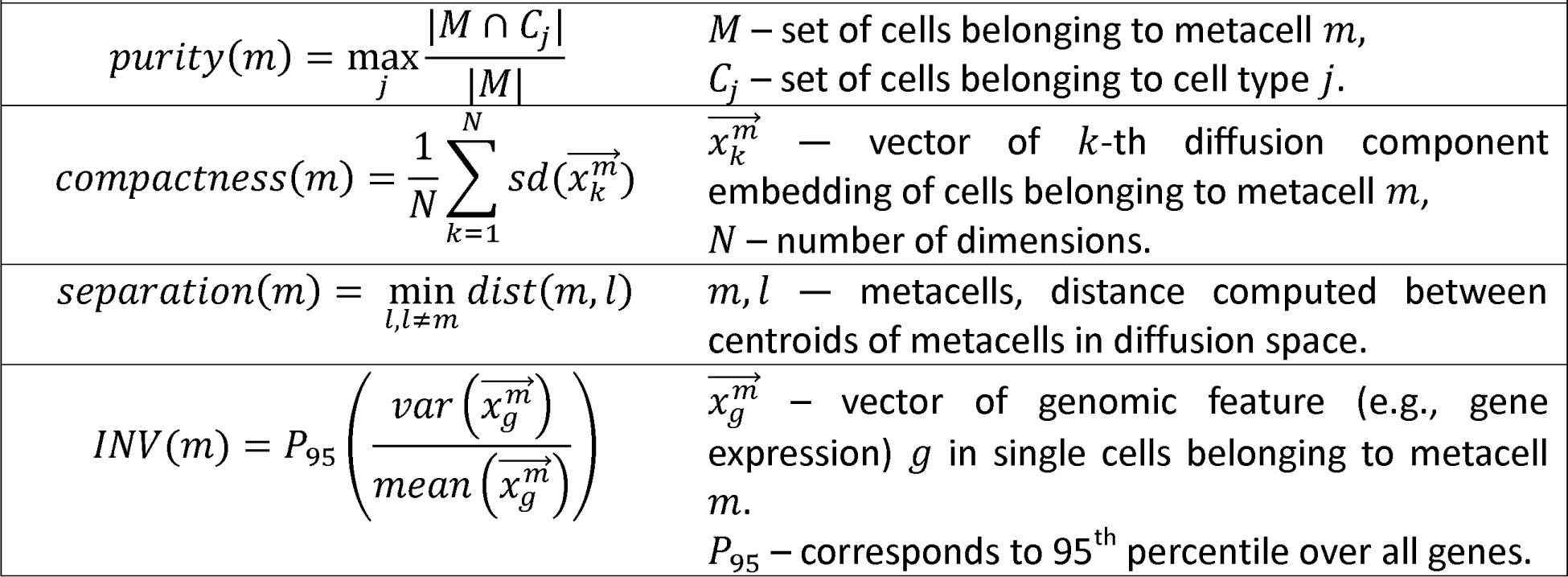
Analytical formulas of metacells quality metrics.

#### Purity

The metacell purity represents the fraction of cells from the most abundant cell type in a metacell (Fig. 3a). In the context of metacells, this measure was introduced in Bilous et al.^40^. Purity is a useful measure to check that metacells do not mix cells from different cell types. Purity requires prior knowledge of cell-type annotation, which is not necessarily available for newly generated datasets. To evaluate metacell purity in a non-annotated dataset, cell-type annotation^71,72^ leveraged from well-annotated reference atlases^73–78^ can be applied to single-cell data.

**Figure 3.**
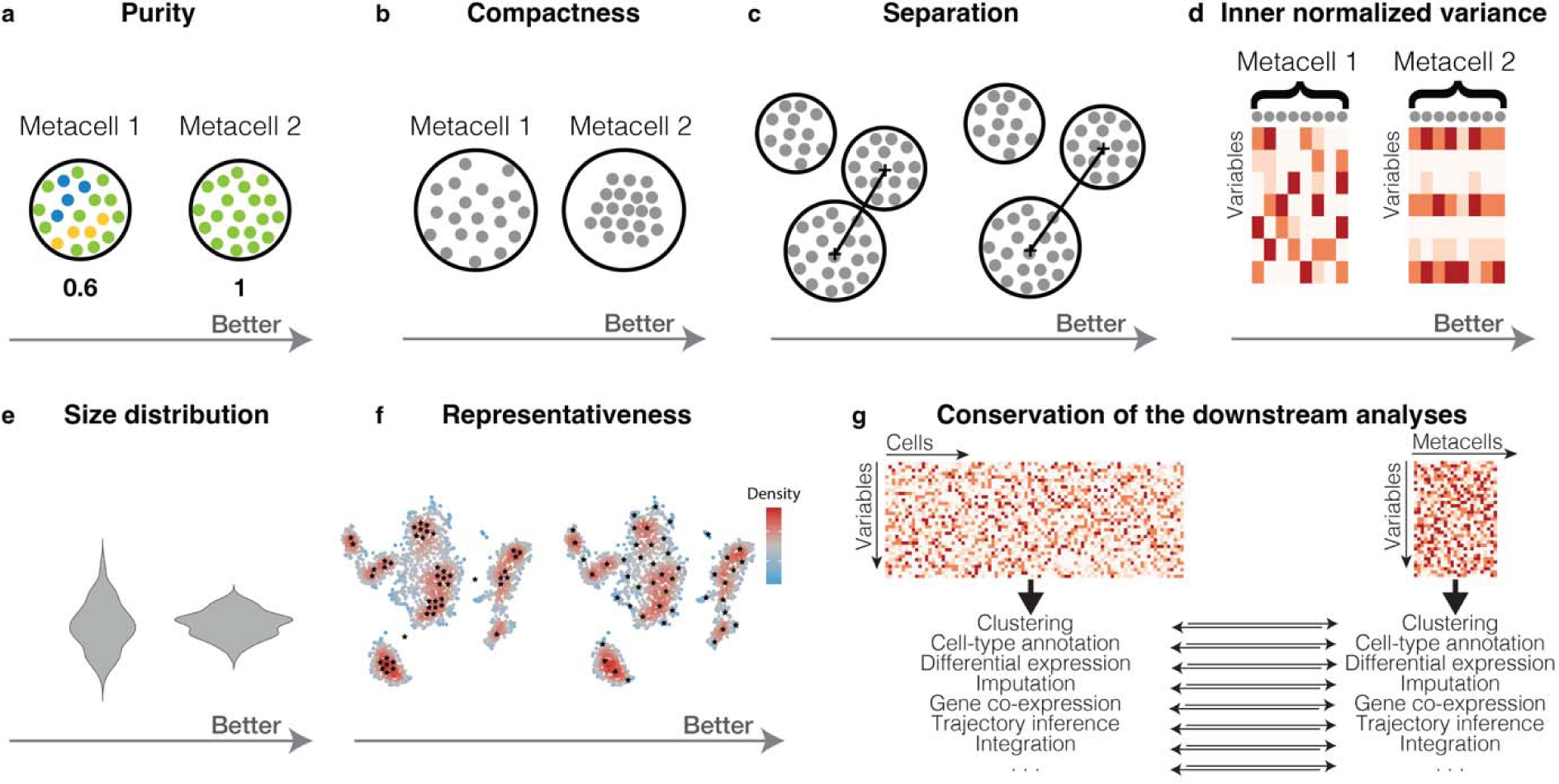
Metacell quality metrics. **a**, Purity is defined as the proportion of cells from the most abundant cell type in a metacell. Higher purity corresponds to higher proportion of cells of the same cell type within a metacell. Purity can also be defined based on other annotations/categories than cell types. **b**, Compactness is defined as the average variance of latent space component within a metacell. Better compactness corresponds to lower variance in the latent space components within cells grouped into a metacell. **c**, Separation is defined as the distance to the closest metacell. Better separation corresponds to more distant metacells in the latent space. **d**, Inner normalized variance is defined as the mean normalized gene variance within a metacell. Better inner normalized variance corresponds to lower variance of the single-cell profiles within a metacell. **e**, Metacell size distribution. Better metacell size distributions have lower dispersion. **f**, Representativeness corresponds to the ability of metacells to faithfully represent the global structure of the single-cell dataset. Better representation corresponds to more uniform coverage of the dataset (black stars represent the centroid of each metacell). **g**, Conservation of the downstream analyses at the metacell level is defined as the ability of metacells to preserve the results of the single-cell analysis.

#### Compactness

The metacell compactness corresponds to its homogeneity and is defined as the average diffusion component standard deviation of single cells within a metacell (Fig. 3b). In the context of metacells, this measure was introduced in Persad et al.^41^. In contrast to purity, compactness does not require prior knowledge of cell-type annotation and therefore can be used to mark potentially low-quality metacells or metacells that need further inspection before any downstream analyses. However, compactness is latent-space dependent, resulting in different values according to the space used to compute it (Supplementary Fig. 1a). For this reason, compactness values are not straightforward to interpret when comparing tools that use different latent spaces to identify metacells. For instance, the MetaCell and MC2 use normalized gene space, while SEACells and SuperCell use the principal component analysis (PCA) space. Therefore, by construction, the latter two will tend to perform better in the metrics computed in the PCA (or a derivative from PCA, such as diffusion components^79^).

#### Separation

The metacell separation represents the remoteness from the closest metacell and is defined as the Euclidean distance between centroids of the single cell coordinates forming the metacells (Fig. 3c). In the context of metacells, this measure was introduced in Persad et al.^41^. Similar to compactness, it does not require prior knowledge on cell-type annotation but is also latent-space dependent (Supplementary Fig. 1b). For those reasons, it can be used for identification of potentially low-quality metacells but is less recommended for comparing metacell construction tools that use different latent space for metacell identification.

There is also a clear relationship between separation and compactness: better compactness results in worse separation (Supplementary Fig. 1c) and vice versa. Metacells from dense regions will have better compactness but worse separation, while metacells from sparse regions will have better separation but worse compactness.

#### Inner normalized variance (INV)

INV corresponds to the mean normalized variance of features (e.g., genes) within a metacell (Fig. 3d). In the context of metacells, this quality measure was introduced in Ben-Kiki et al.^39^. following the idea that the profile of an ideal metacell originates from the same multinomial distribution and therefore should have intrinsic gene variance proportional to its mean. As such, metacells with minimal INV should contain cells that differ mainly due to technical reasons. It is the only metric that does not require prior knowledge and is latent-space independent.

#### Metacell size distribution

The size of a metacell is defined as the number of single cells it contains (or, in rare cases, the total number of UMIs^39^). Statistically, larger metacells tend to express more genes, potentially confounding downstream analyses. To ensure balanced downstream analyses, it is better to have a homogeneous metacell size distribution (Fig. 3e).

#### Representativeness

A good metacell partition should reproduce the overall structure (i.e., manifold) of the single-cell data by uniformly representing its latent space (Fig. 3f). This idea has been used to visually compare different metacell construction approaches^41^. When building metacells, it can be useful to verify their presence across all regions of the single-cell data manifold, including sparse areas. To achieve uniform space coverage, SEACells employs a maximum-minimum approach to determine initial centroids for the future metacells^41,63^.

Akin to a trade-off between compactness and separation, a more uniform representativeness of the manifold leads to increased variation in metacell sizes to compensate for inherent over- and under-representation of different cell types. Abundant cell states can accommodate larger metacells to account for their density, while rare cell types are typically represented by smaller metacells. Therefore, a desirable metacell partition requires to balance those two characteristics.

#### Conservation of the downstream analyses

The metacell concept has been introduced as an alternative representation of single-cell data, offering utility in downstream analyses (Fig. 3g). Therefore, preserving the results obtained at the single-cell level and avoiding artifacts are key criteria for their effective use. Several approaches have been proposed to assess the compatibility of metacells with downstream analysis tools and the conservation of the biologically relevant results obtained at the single-cell level^40^, of which we discuss some.

To evaluate the conservation of clustering results, clustering assignment obtained at the metacell and single-cell levels can be compared using adjusted rand index (ARI)^80^ or adjusted mutual information (AMI)^81^. Preservation of the differential expression analysis can be evaluated by comparing the overlapping significant genes obtained at single-cell and metacell levels. This can be done employing AUC score or true positive rate (TPR), ranking genes by statistical significance (i.e., p-value) and relative deviation in expression (i.e., log fold change). Additionally, the differential expression concordance can be evaluated by comparing functional enrichment of identified signatures^82^, for instance, by comparing the results of gene set enrichment analysis^83^. Pseudotime ordering preservation can be assessed using correlation analysis. For RNA velocity^84^, comparison of angles between the inferred velocity vectors in a joint embedding of the metacells and single-cells has been proposed to quantify the agreement between single-cell and metacells RNA velocity^40^.

The assessment of the conservation of the downstream analyses requires to perform analyses at both the single-cell and metacell levels. This assessment is therefore especially useful for validating newly developed metacells construction or analysis methods, to determine whether their results preserve the biologically relevant information in the single-cell data.

To summarize, high-quality metacells should contain cells of the same cell type, be compact and well separated from other metacells, as well as have low normalized variance. Globally, metacells should decently represent the entire manifold of the single-cell data, while having homogeneous metacell size distribution to ensure that the results of the downstream analyses are driven by biological variation rather than the bias in the amount of information each metacell bares (see section ‘Limitations of metacells’). Several quantitative measures of the quality of metacells can be computed with the metacell quality module provided as an R package in our metacell toolkit repository (https://github.com/GfellerLab/MetacellAnalysisToolkit). Beyond these quantitative quality measures, it is essential that metacells preserve the information present at the single-cell level and do not create artifacts (see section about ‘Limitations of metacells’)^40^.

## Applications of metacells

Metacells have found extensive applications in conducting downstream analyses of single-cell genomics data. Such analyses were performed for various types of transcriptomics data from diverse biological context, including hematopoiesis^85^, tumorogenesis^86^, cancer immunology^51,87,88^, host-viral interaction^89^ organ development^90^ and whole-organism profiling^91^. Below we describe some of the standard analysis that can be performed with metacells.

### Data normalization

Sparsity of single-cell data poses challenges to every step of the downstream analyses. This also applies to the data pre-processing, where normalization is used to counter cell-specific biases. To address this, the idea of pooling similar cells for size factor estimation^92^ was proposed even before the metacell concept was introduced.

### Visualization

Metacells have been used for more compact data visualization in multiple studies^51,87,89–91,93,94^. Different approaches for displaying metacells in a reduced space have been proposed. When complementing single-cell visualization, metacells have been overlaid onto the low-dimensional representation of single cells by positioning each metacell at the centroid of the single cells it comprises^38,41^. Alternatively, metacells can be visualized independently by applying dimensionality reduction tools (e.g., PCA, tSNE^64^, UMAP^95^) on the metacell profile matrix^39–41^. Although most of the studies display metacells as entities of equal size, the variability of metacell sizes can be visualized using shapes of proportional sizes. We and others have observed that in general, the visualization performed directly on metacell profiles preserves most of the patterns observed in the single-cell data^39,40^.

### Clustering

Unsupervised clustering is commonly applied to single-cell data to identify cell types and cell states. Similarly, clustering was applied to metacells when performing downstream analysis at the metacell level^51,94^. In our work, we observed that most clusters identified at the metacell level recapitulate those observed at the single-cell level, and differences fall within the range of fluctuation observed by using different clustering algorithms^40^.

### Differential expression

Differential expression in scRNA-seq data aims at identifying signature genes by comparing the level of expression of a gene between two groups of cells. When performing downstream analyses at the metacell level, differential expression was performed between groups of metacells^51,94^. Differential expression analysis at the metacell level, either between conditions or between clusters, has been shown to recapitulate most results observed in single-cell data^40^.

### Cell-type annotation

Metacells can be annotated to a particular cell type in different ways, using: (i) clustering-based, (ii) metacell-based or, (iii) single-cell-based approaches. The first approach relies on metacell clustering and differential expression, followed by analysis of the differentially expressed gene in each metacell cluster. This approach integrates the overall structure of the data in the annotation but depends on the accuracy of the clustering algorithm. The second approach annotates each metacell to a specific cell type based on known marker genes, similar to many single-cell annotation pipelines^71,72^. The third approach (single-cell-based) assumes that a single-cell annotation is available and annotates a given metacell to the most abundant cell type within this metacell.

### Gene co-expression analyses and gene regulatory network inference

Since metacells have less sparse profiles, they have been used for signature enrichment and gene co-expression analysis^85,94,96–98^. Gene co-expression, which is used for gene regulatory network (GRN) analysis, often suffers from the high dropout in single-cell data. For this reason, an aggregation of profiles of highly similar cells for GRN analysis was used even before the introduction of the metacell concept^99^. Currently, the metacell concept is used to enhance signal for GRN construction for scRNA-seq^100–103^, for more sparse single-cell lncRNA-seq data^104,105^ and for multi-omics data^106^. Given the computational cost of GRN analysis^17^, metacells facilitate scaling of GRN construction tools, particularly those initially developed for bulk RNA-seq^48^, to handle larger datasets^100^.

### Trajectory analyses

Metacells have the potential to facilitate more advanced steps of single-cell analysis such as inferring developmental trajectories^40,41,56^. In our own work, we observed that RNA velocity^84^ inferred at the metacell level provides highly similar results as those observed in single-cell data^40^. It has been also demonstrated that pseudotime analysis at the metacell level recapitulates known developmental dynamics^41^, although the conservation of pseudotime between single cells and metacells has not yet been formally demonstrated.

### Data integration and atlas construction

Data integration represents a computationally demanding yet indispensable step in single-cell data analysis, considering the abundance of datasets produced from various technologies^2,20–22,107,108^ and the high interest in constructing single-cell atlases of organs or even organisms^73–78^. The use of metacells prior to data integration has demonstrated several benefits. First, it streamlines the integration process, enabling the integration of millions of cells on a standard laptop^40^. Second, it has been shown to enhance integration results by revealing hidden cell populations that may remain undetected when directly integrating single-cell data^41,51^. This improvement is attributed to the fact that integration tools often overcorrect for variability within and across samples. By reducing excessive within-sample variability, metacells have been shown to help integration algorithms to avoid overcorrection^41^.

Improved integration results, along with reduced computational burden and enhanced biological signal, strongly support the use of metacells for the construction, analysis, and storage of single-cell atlases^109^. For instance, Tumor Immune Single-cell Hub 2 (TISCH2)^78^ utilizes metacells (referred to as mini-clusters in the original study) to perform computationally expensive gene-gene correlation analysis in a single-cell atlas comprising over 6 million cells.

### Accelerating downstream analyses

Many computational tools for single-cell analysis are not scalable to current large-scale datasets, particularly those that were initially developed for the analysis of bulk RNA-seq data. One of the main purposes of using metacells is to decrease the computational burden associated with the large size of single-cell genomics datasets, both in terms of time and memory usage. Given that each step of the downstream analyses is computationally demanding, metacells significantly accelerate each of them and the overall analysis^40^. For this reason, many tools use^110^ or suggest to use^111,112^ metacell approaches to scale their proposed methods to large datasets. These include also methods for cross-modality cell matching^110^, metabolic state inference^111^, and data integration^112^.

Overall, metacells have been frequently applied for conventional single-cell downstream analyses, enabling efficient handling of large datasets and scaling tools initially designed for bulk RNA-seq. For this reason, many single-cell analysis tools adopt metacell-like approaches to handle large datasets.

Some studies, including ours, not only reported conservation of the results of analyses performed at the single-cell level, but also improvements based on metacells (i.e., biological signals that are difficult to capture in single-cell data are more visible in metacells)^40,41,51^. While this is true in some cases, it is still important to realize that, by construction, metacells are a direct aggregation of single cells. Therefore, any biological information observed in metacells is likely to be present, though possibly more noisy and less visible, in the single-cell data. Cases where metacells would reveal patterns that are totally absent or invisible in the single-cell data, should therefore be treated with care (see also discussion in the ‘Limitations of metacells’ section).

## Metacells as a trade-off between sketching and imputation

By aggregating information within several highly similar cells, metacells reduce the size of the dataset and amplify the biological signal. This simultaneously addresses two main challenges of single-cell genomics data analysis: the large size of the single-cell data^1^ and its excessive sparsity^34^. Individually, these challenges have been addressed by sketching (i.e., topology-preserving downsampling)^24,26^ and data imputation^27–33^.

Sketching is a powerful way to reduce the complexity of large datasets, and optimized approaches have been developed to handle rare cell types or sparsely sampled regions of the single-cell landscape. A major advantage of sketching is the limited risk of creating artifacts due to the aggregation of cells that are biologically distinct (e.g., impure metacells). However, by overlooking a large fraction of the cells, one cannot exclude that the reduced data will lose some of the information present in the initial data. Moreover, the reduced data suffer from similar sparsity as the original data.

Imputation capitalizes on cells that share some transcriptomic similarity to refine the profile of each cell based on the ones of its neighbors. As such, the imputed data become much less sparse and can reveal biological patterns difficult to capture at the single-cell level^29–31^. Unfortunately, the imputed data are even larger than the initial one, and imputation has been shown to introduce false signals in data^113^.

Thus, metacells emerge as a trade-off between sketching and imputation by simultaneously addressing the challenges posed by the large-scale nature and high sparsity of single-cell data (Fig. 4).

**Figure 4.**
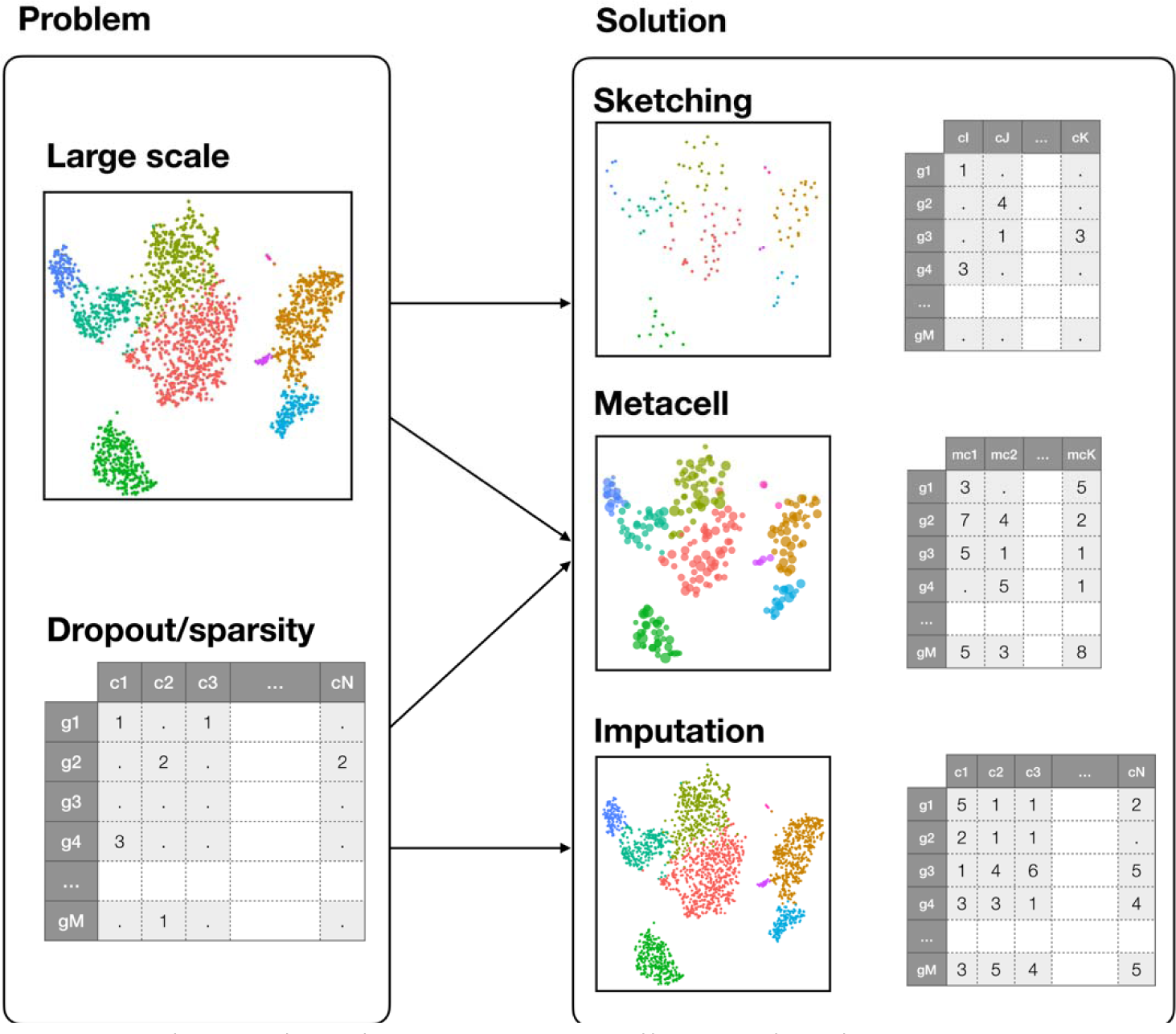
Relationships between metacells and sketching or imputation. Metacells combine the reduction in size of sketching approaches and the reduction in sparsity of imputation strategies.

## Limitations of metacells

Metacells have been successfully used in several studies, ranging from data visualization to downstream analysis and method scaling^51,85,87,89–91^. Provided a reasonable graining level is used, we and others have observed that metacells capture most of the biological information present in the single-cell data and preserve the results of downstream analyses. Still, as with all data transformation approaches, artifacts can appear with metacells. Below, we highlight some of these issues and recommend strategies for mitigating them.

The metacell partition may be considered a very high-resolution clustering. As with any unsupervised clustering, it can potentially group cells of distinct types within a single metacell (Fig. 5a), resulting in the formation of impure metacells. The aggregated profile of such metacells may represent an artifact. Such artifacts can lead to misleading interpretations, including the presence of non-existing intermediate states or spurious gene co-expression (Fig. 5a). Additionally, rare cell types could be completely missed if entirely aggregated with a more abundant cell type into a single metacell.

**Figure 5.**
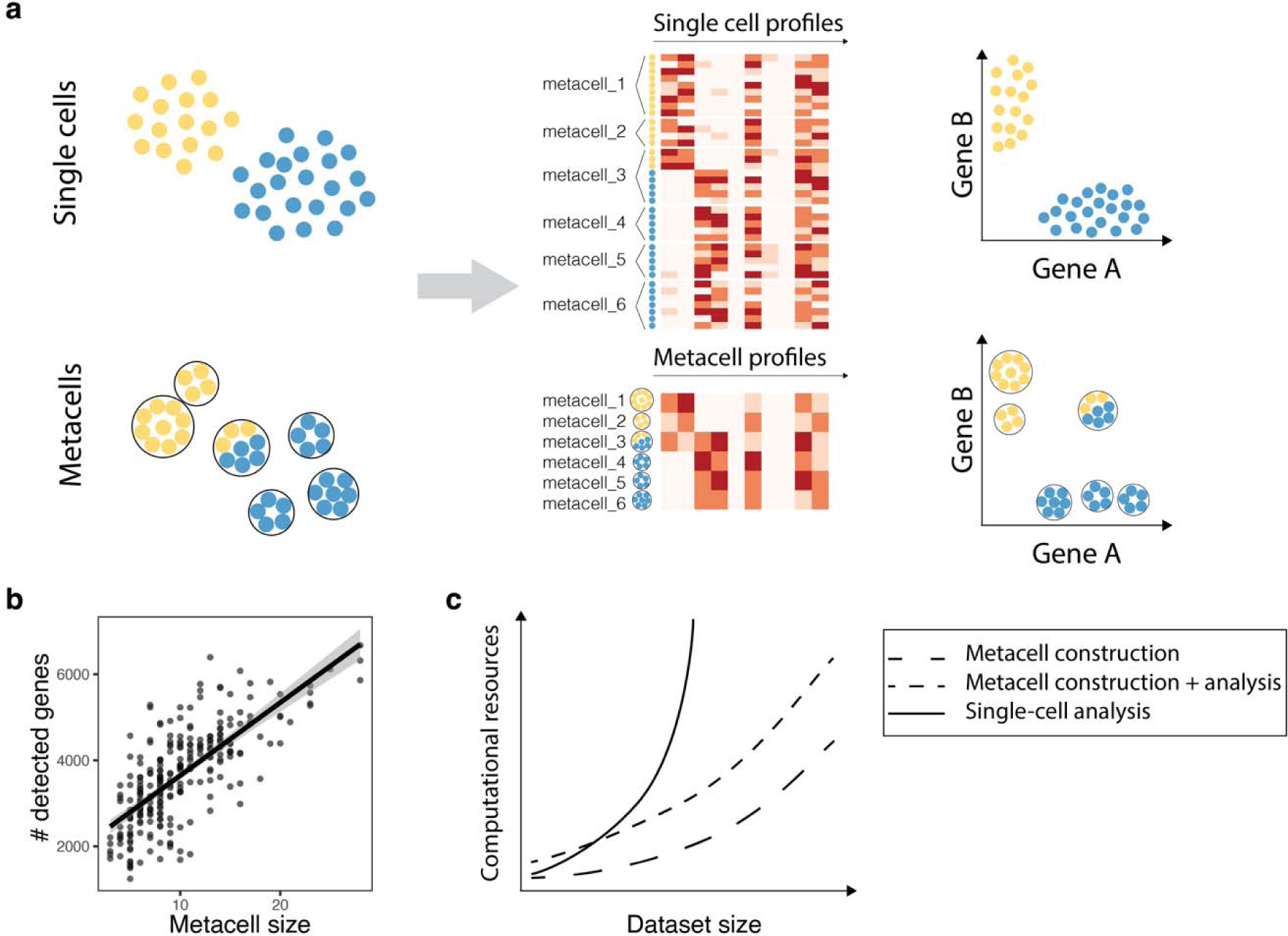
Limitations of metacells. **a**, Example of limitations in metacells when aggregating cells of different cell types (i.e., impure metacell_3 in the example). Such impure metacells can lead to mixed profiles and artifacts in gene co-expression analyses. **b**, Correlation between the size of metacells and the number of detected genes. **c**, Illustration of the computational cost of metacell construction, metacell construction + analysis, and single-cell analysis.

A first approach to tackle this issue involves constructing metacells at a lower graining level, thereby achieving a higher resolution. An alternative way to prevent aggregation of cells from different cell types is to build metacells in a supervised manner by constructing metacells for each cell type separately^86^ or by splitting impure metacells^40^ when an annotation of the single cells is available.

Another limitation specific to metacell analysis is the skewness in the distribution of metacell sizes, which arises from the uneven density and heterogeneity of phenotypes. The discrepancy in metacell sizes leads to differences in profile coverage, with larger metacells having more detected genes (Fig. 5b). Consequently, some observations at the metacell level may be driven by the metacell size. To address this issue, strategies such as balancing metacell size distribution and considering metacell size information in downstream analyses can be employed. For instance, to avoid very large outliers, the MetaCell approach dissolves metacells exciding a particular size^38,39^. Alternatively, the SuperCell tool provides the analytical framework that accounts for metacell size in downstream analyses. Our benchmarking^40^ suggests that considering metacell sizes in metacell analyses is especially important for higher graining levels (50-100).

Finally, metacells construction can be computationally demanding since many existing tools require computing PCA and/or building a similarity network between cells. Several approaches have been developed to accelerate the metacell construction procedure. One approach adopts a divide-and-conquer methodology^39^ by randomly splitting large datasets into piles and performing metacell construction within them. Another approach constructs metacells for a subset of cells and projects the remaining cells onto the constructed metacells^40^. Topology-preserving downsampling (i.e., sketching)^24,26^ should improve the representativeness of such an approach. A third approach involves constructing metacells per sample or other identities with a consecutive integration of the resulting metacells (see ‘Workflow recommendations’ section). In general and especially for large datasets, the downstream analysis at the metacell level shows a significant acceleration (Fig. 5c), effectively compensating for the metacell construction time^40,41^, especially for exploratory analyses where many tools or choices of parameters are tested on the same data.

## Concepts that share similarities with metacells

Some of the ideas underlying the metacell concept, such as aggregation of profiles and/or representation of data at different resolution(s), have been used in other approaches. These include nested communities, graph abstraction, neighborhoods, pseudobulks, pseudocells and pseudoreplicates (see Box 2).

#### Box 2. Concepts that share similarities with metacells.

**Nested communities** or **pan-resolution representation** – representation of data in a hierarchical manner^44,114,115^, wherein high resolution can be seen as a metacell partition.

**Graph abstraction**^116^ – representation of single-cell data as a graph of clusters at different resolutions (cell types or subtypes) for further investigation of cell-type relationships.

**Neighborhood** – partition of the single-cell data into partially overlapping clusters of transcriptionally highly similar cells. Neighborhoods can be seen as partially overlapping metacells. Neighborhood can be defined for all single cells^27,117^ or for a set of anchoring cells^118,119^.

**Pseudobulk** – partition of cells based on metadata (e.g., samples, conditions) and/or broad transcriptomic similarity (i.e., cell types), followed by the aggregation of their profiles.

**Pseudocell** – aggregation of profiles of randomly sampled cells (potentially with replacement) to increase sample size^120^.

**Pseudobulk of pseudoreplicate** – random assignment of cells of one replicate to several artificial replicates followed by aggregation of their profiles^121^.

*Nested communities* – represented by tools such as TooManyCells^114^, multiscale PHATE^44^, and HiDeF^115^ – enable exploration of single-cell data at multiple resolutions by providing its hierarchical structure (Fig. 6a). Usually, the root corresponds to the entire dataset branching into types and sub-types. A representation at a high resolution may be considered as a partition of single-cell data into metacells.

**Figure 6.**
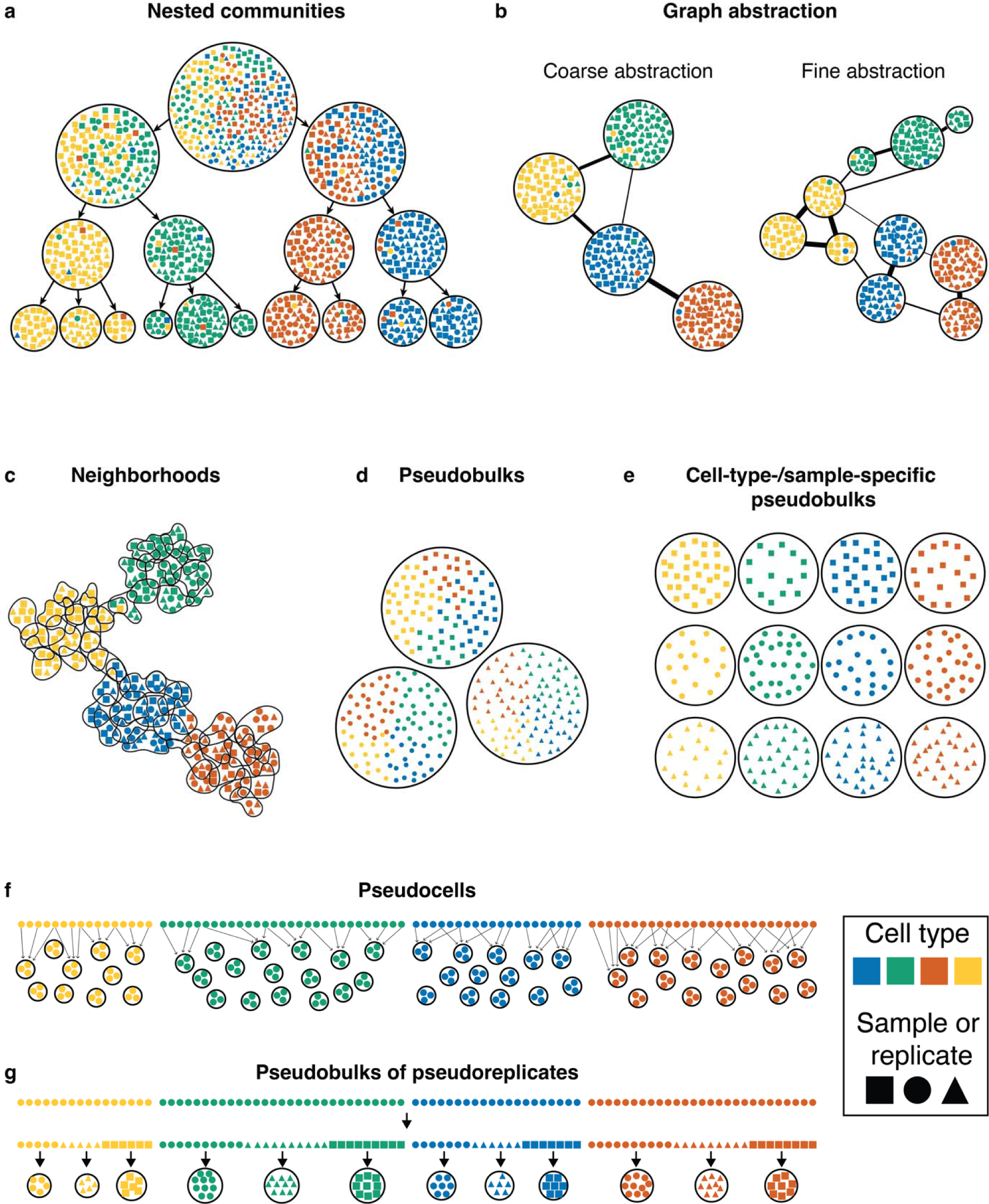
Concepts that share similarities with metacells. **a**, Example of nested communities. **b**, Example of graph abstraction. **c**, Example of neighborhoods. **d**, Examples of sample-specific pseudobulks. **e**, Example of cell-type-/sample-specific pseudobulks. **f**, Example of pseudocells. **g**, Example of pseudobulks of pseudoreplicates.

*Graph abstraction*^116^ is a representation of single-cell data as a graph of cell types (Fig. 6b) to study relationships among them, including putative trajectories. It also allows users to investigate data at different resolutions by providing graph abstraction at multiple graining levels. In contrast to metacells, graph nodes represent cell types or clusters, rather than groups of highly similar and ideally redundant cells.

*Neighborhood* is a partition of data into partially overlapping groups of highly similar cells (Fig. 6c). The concept of neighborhood is very similar to the one of metacell, except that the neighborhoods partially overlap, and cell profiles are not necessarily aggregated^119^.

*Pseudobulks* represent an aggregation of cells based on metadata, cell-type annotation or clusters. Specifically, aggregations of cells based on sample identity correspond to the sample-specific pseudobulks (Fig. 6d), and aggregations of cells based on both samples and cell types correspond to cell-type-/sample-specific pseudobulks (Fig. 6e).

*Pseudocells* are constructed by aggregating randomly sampled cells from the same cell states, potentially with overlaps (Fig. 6f). This allows to compensate for low number of cells and high sparsity^120,122^. *Pseudoreplicates* are generated by randomly assigning cells from one replicate to several artificial replicates. *Pseudobulk of pseudoreplicates* ^121^ are obtained upon aggregating pseudoreplicates (Fig. 6g).

These different approaches share similarities with metacells but correspond to different concepts. In rare cases, the term ‘metacell’ was used when referring to some of these approaches, like neighborhood^118^, pseudocell^17^ or pseudobulk^123,124^. To avoid ambiguities, we recommend using the term ‘metacell’ only for disjoint sets of highly similar cells whose profiles are aggregated.

## Workflow recommendations

Metacell construction and analysis involve several steps that have been discussed before. To guide users interested in building and analyzing metacells from their own data, we provide recommendations for all essential steps of the process. We further refer users to our tutorial (https://github.com/GfellerLab/MetacellAnalysisTutorial) and toolkit (https://github.com/GfellerLab/MetacellAnalysisToolkit), where code is provided for most of these steps.

### 1. Metacell construction

As with any single-cell genomics analysis, the metacell construction starts with a single-cell profile matrix. Since the metacell concept was initially introduced for single-cell transcriptomics data, here we assume that the single-cell profile matrix is a cell-by-gene matrix. We also assume that the low-quality cells and genes are filtered out and profiles are library size-normalized and transformed (see refs.^125,59,126,127^ for recommendations). To build metacells, the following steps should be performed on the normalized counts matrix:

#### 1.1. Feature (or highly variable genes) selection

Given the high dimensionality of single-cell genomics data (e.g., ∼20’000 genes), it is recommended to first select the most informative genes (typically the top 2’000 most variable genes). Certain types of genes, like mitochondrial, ribosomal, and stress-related genes, can also be left out from the analysis as their expression is often influenced more by technical factors than actual biological distinctions^128^. All the available single-cell analysis frameworks provide implementations of several common approaches for feature selection, and the construction of metacell is not restricted to a specific one (except for the MetaCell and MC2 tools, which relies on the in-built approach for the selection of informative genes, thus does not require this step).

#### 1.2. Dimensionality reduction (usually PCA for transcriptomics data)

To quantify distances or similarities between cells, we recommend using dimensionality reduction approaches suitable for a particular modality (e.g., PCA for transcriptomics). The number of principal components can be chosen the same way as for most other scRNA-seq data analyses (typically between 10 and 50). This step is not necessary for the metacell construction with the MetaCell and MC2, as these tools build metacells from gene counts space relying on in-built data transformations.

#### 1.3. Partitioning data into metacells

For metacell construction with Metacell and MC2, raw counts should be used. For metacell construction with SuperCell, either normalized counts or low-dimensionality representation should be used. For metacell construction with SEACells, low-dimensional representation should be used. As an alternative to these three methods, metacells can be computed as an excessive clustering of data with a user-defined similarity metric.

#### 1.4. Choice of the graining level

When using tools that allow to pre-decide the graining level (i.e., level of size reduction between the single-cell and the metacell data) we recommend selecting graining levels between 20 and 75. This can be done easily in SuperCell, SEACells, and in the latest version of MC2 (v 0.9.3). In MetaCell and the original version of MC2, the final graining level is determined by the choice of the minimum or target size of metacells (measured in number of single cells or total number of UMIs, respectively). Therefore, it may happen that the final graining level is far from the expected one or that the resulting metacell partition is too coarse or too fine. To obtain a finer resolution, the minimum or target size of metacells has to be decreased. Most methods require re-running the metacell partition for each choice of the graining level. Exceptions include tools that provide a hierarchical structure of the metacell partitions (SuperCell or other hierarchical clustering algorithms), allowing for fast scanning across multiple graining levels.

To choose an appropriate graining level for a particular dataset, we recommend considering both the size and the complexity of the dataset (see Fig. 2d,e). To preserve the statistical power of the metacell downstream analyses, we suggest using graining levels such that the resulting number of metacells is at least ten times larger than the expected number of fine-resolution cell types. We do not recommend using metrics developed to find the optimal number of clusters, such as modularity^65^, silhouette coefficient^66^ or other metrics developed for scRNA-seq data^67–69^ to determine the number of metacells.

#### 1.5. Note on metacell construction for multi-sample datasets

If a dataset contains several samples and/or conditions, the metacell partition of such data can be done in two ways. The first one is to construct metacells across all samples and conditions, allowing to mix samples and conditions within one metacell^85,87,89,98,129–132^. The second one is to construct metacells per sample and/or per condition, obtaining sample- and/or condition-specific metacells^51,52,78,86,94,123,133,134^. The choice of the method depends on the specific objectives of the downstream analysis and size of samples/conditions. The first approach – cross-samples and conditions – is suitable for analyzing global heterogeneity in the dataset and enrichment of conditions within specific cell states (i.e., metacells). In this approach, the batch effect must be corrected before the metacell construction. Considering that metacells are computed from the entire dataset, the first approach is more compatible with smaller samples. The second approach – per-sample and/or condition – prevents metacells consisting of cells from multiple origins and is more appropriate for conducting differential expression and differential abundance analysis among conditions, as well as for data integration. Samples have to be large enough to have sufficient redundancy to build pure metacells.

#### 1.6. Note on metacell construction for annotated datasets

In the presence of reliable annotation of the single-cell data, it can be useful to integrate this information in the construction of metacells. This can be done in two ways. The first one involves creating metacells per cell type, akin to per-sample metacell construction in multi-sample datasets. The second approach is to split metacells containing mixed cell types after metacell partitioning. We recommend the first approach, as it enables the formation of more specific metacells through the use of cell-type-specific informative features. Additionally, the second approach may lead to lower graining levels and more heterogeneous distribution of metacell sizes due to an increased number of singletons.

#### 1.7. Metacell profile aggregation strategy

When obtaining metacell profiles, single-cell profiles are aggregated using different methods: MC2 computes gene fractions, SuperCell averages normalized counts by default (with an option to sum raw counts), and SEACells sums raw counts. We recommend summing raw counts for comprehensive downstream analyses, as many downstream analysis methods operate on raw counts or require particular normalization^135–137^. The averaging of normalized counts approach is suitable when metacells serve a specific task like visualization, annotation, co-expression analysis, gene regulatory network inference, or trajectory analysis as it does not require re-computing all the upstream steps starting from scaling and normalization.

#### 1.8. Visualization

To visualize metacells, we recommend two approaches. The first one involves overlaying metacells with single-cell data by placing each metacell at the centroid of its constituent cells in a low-dimensional space, which is useful for comparing metacells with single-cell data. It is also possible to combine metacells and single cells to create a joint low-dimensional space. The second approach visualizes only metacells, deriving a low-dimensional representation directly from the metacell profiles. This approach is recommended for exploring the structure of single-cell data at the metacell level for very large datasets (e.g., >1M cells) where visualizing single cells can become computationally heavy.

To facilitate the exploration of these different steps in metacell construction, a practical guide comprising the MC2, SuperCell, and SEACells methodologies can be found in the tutorial accompanying this review (https://github.com/GfellerLab/MetacellAnalysisTutorial).

### 2. Downstream analyses of metacells

Metacells are meant to streamline the analysis of single-cell genomics data. Most of the studies analyze metacells the same way as single-cell data^51,85,87–90,97,98,138,139^ or suggest doing so^39,41,44^. However, metacells have different sizes and thus carry different amounts of information. Therefore, we suggest considering tools that account for sample weight (i.e., metacell size), in particular for analyses that are based on comparing relative values (e.g., differential expression and differential abundance analyses). Below, we discuss some of the most common analyses performed on metacells.

#### 2.1. Pre-processing

Before running downstream analyses, we encourage users to verify that metacells are of good quality. Additionally, the metacell profiles have to be scaled and normalized, especially if the metacell profile corresponds to a sum of counts.

#### 2.2. Feature selection and dimensionality reduction

In general, and especially for unsupervised analyses, we recommend using the same set of features (i.e., informative genes) as the one used for metacell construction. In situations involving multiple feature sets (e.g., per-sample metacell construction) and therefore requiring data integration, we recommend recomputing common features at the metacell level.

Low dimensional embedding (e.g., PCA and further UMAP) can be performed at the metacell level using a set of features defined above.

#### 2.3. Clustering

Any clustering algorithm can be applied on metacells. However, we recommend using clustering approaches that consider sample weight (i.e., metacell size), such as hierarchical clustering or partitioning around medoids^140,141^, especially for large graining levels or for metacells with very heterogeneous size distribution.

#### 2.4. Differential expression

Metacells are compatible with most differential expression methods. However, we recommend using methods that account for weights (e.g., weighted t-test), since neglecting metacell size can lead to erroneous group average gene expression estimates, impacting differential expression analysis results (Fig. 7b).

**Figure 7.**
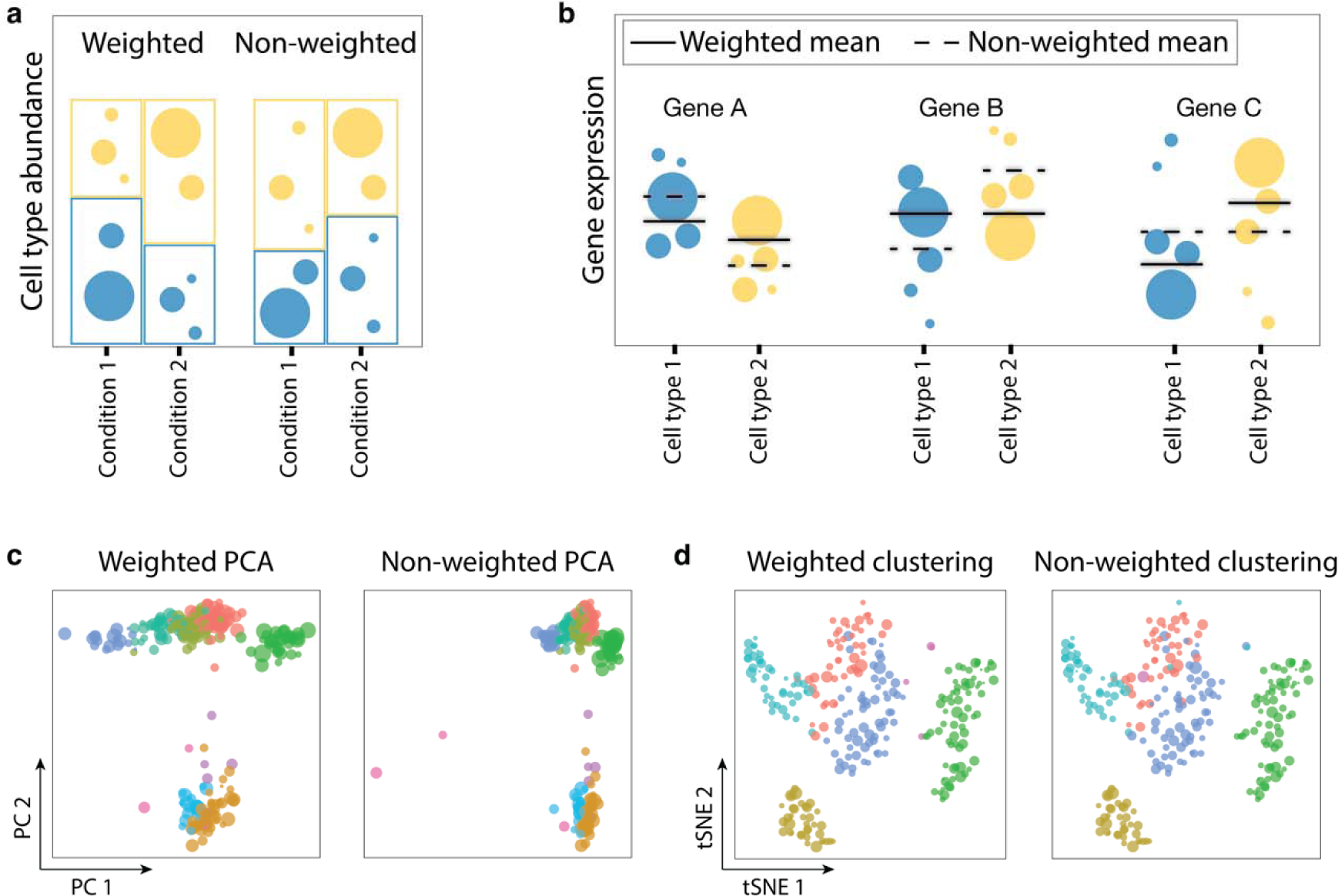
Impact of metacell sizes on the results of the downstream analyses. **a**, Comparison of the results of weighted versus non-weighted differential abundance analysis at the metacell level. Each dot is a metacell colored by cell type. Bars correspond to the estimated proportions of each cell type in a condition with and without considering the size of each metacell. **b**, Comparison of the results of weighted versus non-weighted differential expression analysis. Each dot is a metacell colored by cell type. Solid and dashed lines correspond to weighed and non-weighted estimation of mean expression. **c**, Results of weighted and non-weighted principal components analysis for the same dataset. Each dot is a metacell colored by cell type. Better separation of cell types is observed in the weighted PCA. **d**, Results of weighted and non-weighted Louvain clustering, with dots representing metacells colored by cluster annotation. Size of dots correspond to the size of metacells.

#### 2.5. Metacell annotation

When a reliable annotation of the single-cell data is available, we recommend using it for metacell annotation. This further allows users to check the purity of metacells. In the absence of single-cell annotation, we advise metacell annotation based on markers, similar to single-cell data. In our experience, improved gene coverage in metacells leads to their better annotation compared to single-cell data^40^. We also recommend comparing this metacell-based annotations with metacell clustering/visualization results, akin to single-cell data annotation practices.

#### 2.6. Gene co-expression

Metacells, by aggregating information from multiple cells, offer a more comprehensive profile with reduced dropout rates, making them more suitable for gene co-expression analysis. To enhance the accuracy of these analyses, we recommend using weighted correlation methods that consider the metacell sizes, ensuring a more precise assessment of gene co-expression patterns. In drawing scientific conclusions, it is essential to verify that reported gene co-expression is consistently present at the single-cell level (although at lower significance) to make sure it does not come from potential metacell artifacts (Fig. 5a).

#### 2.7. Velocity

Metacells are compatible with RNA velocity^84^ analysis for inferring potential developmental trajectories. Metacells can be computed from all counts or spliced counts only. The standard RNA velocity algorithms use a neighborhood approach to impute data for estimating an equilibrium parameter. To prevent over-smoothing, we recommend using lower graining levels when constructing metacells or lower neighborhood size parameter when running RNA velocity on metacells.

#### 2.8. Note on the use of metacell weights in downstream analyses

An important aspect to consider when analyzing data at the metacell level is the size of each metacell. For instance, when computing cell-type abundance between two conditions, just counting metacells can lead to inaccurate estimations (Fig. 7a). Disregarding the size of metacells when comparing the mean expression of a gene between conditions can create similar artifacts (Fig. 7b), which may lead to type I (Fig. 7b, Gene B) or type II (Fig. 7b, Gene C) errors. In general, it is therefore advised to consider the size of metacells and use downstream analysis tools which are compatible with sample-weighted data. Although weights do not necessarily result in dramatic changes in all analyses (see examples in Fig. 7c-d), our recent work suggests that weights do improve most results for larger graining levels^40^, particularly in light of the accumulation of discrepancies during downstream analysis steps.

More examples of the usage of metacells in the downstream analyses are available in our tutorial (https://github.com/GfellerLab/MetacellAnalysisTutorial).

## Conclusion and outlook

Metacells aim at preserving and possibly improving biological signals in single-cell genomics datasets while reducing their size to facilitate downstream analyses. Many tools, such as MetaCell^38^, MC2^39^, SuperCell^40^ and SEACells^41^, are available to build metacells. Metacells have been effectively employed in various studies for multiple types of downstream analyses^51,85,87,88,90,100–105,139^, and have demonstrated utility in the analysis and representation of single-cell atlases^51,78,134^. Moreover, metacells have been used to scale existing computational methods to larger datasets^110–112^. Performing data integration at the metacell level has been shown to not only facilitate the process^40^ but also enhance the results by diminishing within-sample noise^41,51^. In the future, we anticipate that metacells will be especially useful for the construction, analysis and storage of very large single-cell atlases^142^. Additionally, a synergy between sketching^24,26^ and metacell is expected, where sketching provides a comprehensive low-scale representation complemented by enhanced profiles obtained through metacells.

As with any data transformation approach, metacell can introduce artifacts, especially at large graining levels. In our experience, most of these artifacts arise when grouping transcriptionally similar cells from different cell types (e.g., mixing CD8 and CD4 T cells within the same metacell), or when disregarding the fact that metacells can have different sizes. Being aware of such potential artifacts is therefore important when using and analyzing metacells. To this end, we recommend always evaluating the purity of metacells based on some cell type annotations (either experimentally available or based on prediction tools). We also recommend using downstream analysis pipelines that can account for metacell sizes (i.e., weights).

As of today, metacells have been mainly applied in single-cell transcriptomics, but this concept can be extended to other modalities^41,44^ and to multimodal data^45^. Considering that such data are increasingly being generated at large scales, we anticipate that metacell construction pipelines will be used and tailored for them.

In summary, metacells significantly increase profile coverage (i.e., decrease sparsity due to transcript dropout in scRNA-seq data) and reduce computational resources needed for downstream analyses while preserving the biologically relevant heterogeneity (Fig. 8). As such, metacells can be seen as an optimized structure between the sparse and partially redundant level of single cells and the over-simplified level of clusters. Provided that limitations are well understood and reasonable graining levels are used, metacells offer us a powerful framework for visualizing, analyzing, storing and sharing large-scale single-cell genomics data.

**Figure 8.**
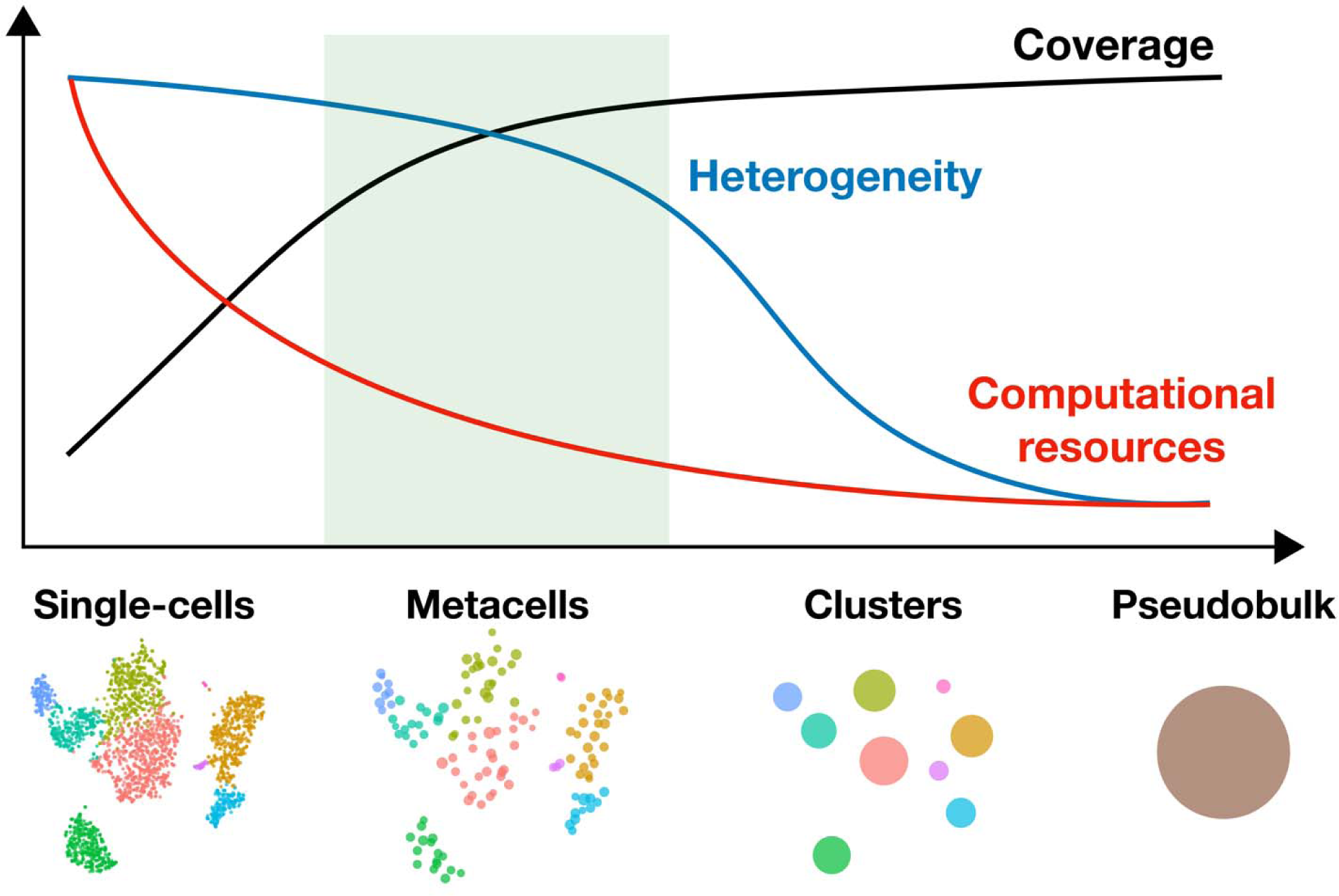
Metacells improve profile coverage and save computational resources, while preserving biological heterogeneity in single-cell genomics data.

## List of abbreviations

ARI: Adjusted Rand Index
AMI: Adjusted Mutual Information
AIR: Adaptive Immune Receptor
AUC: Area Under the Receiver Operating Characteristic Curve
BCR: B-cell receptor
CITE-seq: Cellular Indexing of Transcriptomes and Epitopes by Sequencing
CyTOF: Cytometry by Time of Flight
FACS: Fluorescence-Activated Cell Sorting
GRN: Gene Regulatory Network
INV: Inner normalized variance
kNN: k-Nearest Neighbor
lncRNA-seq: Long non-coding RNA sequencing
PCA: Principal Component Analysis
RNA: Ribonucleic Acid
scATAC-seq: Single-cell Assay for Transposase-Accessible Chromatin using sequencing
scRNA-seq: Single-cell RNA sequencing
TCR: T-cell Receptor
TPR: True Positive Rate
tSNE: t-distributed Stochastic Neighbor Embedding
UMAP: Uniform Manifold Approximation and Projection
UMI: Unique Molecular Identifier

## Materials and Methods

### Dataset for illustrating single-cell, metacell, sketching and clustering representations (Fig. 2a,d; Fig.3f, Fig. 4; Fig. 7c-d; Fig. 8)

PBMC data from 10X Genomics downloaded from the SeuratData package^143^ was used for the majority of illustrative (Fig. 2a,d; Fig. 3f, Fig. 4; Fig. 7c-d; Fig. 8) and analytical (Fig. 5b; Supplementary Fig. 1) purposes. A standard downstream analysis pipeline was applied to obtain a single-cell tSNE^64^ representation of data. Cells are colored according to the provided cell-type annotation. Metacell representation was obtained upon applying the SuperCell approach. A 2D representation of metacells was computed by averaging tSNE coordinates of single cells within each metacell (Fig. 2a, Fig. 4, Fig. 7d, Fig.8). 2D representation of clusters (cell types) was obtained by averaging the tSNE coordinates of single cells within each cluster (Fig. 2a; Fig. 8). The size of metacells, resp. clusters, in tSNE is proportional to the number of single cells in metacells, resp. clusters.

### Computing compactness and separation from different latent spaces (Supplementary Fig. 1)

To compute compactness and separation for different metacell construction tools from different latent spaces (Supplementary Fig. 1), PMBC dataset from 10x Genomics was used as it is, without filtering cells or genes. For MC2 (metacells v.0.8.0), the divide_and_conquer_pipeline() was used with the target_metacell_size being 76’100 UMIs (to obtain the requested graining level of 30). For SuperCell (v.0.1), we used the 10 principal components computed from the top 1000 variable genes with a graining level of 30. The same PCA embedding and graining level was used for SEACells (v.0.2.0) by requesting 87 metacells, initializing algorithm considering 10 eigenvalues and fitting 25 iterations, with the convergence tolerance of 1e-5. Compactness and separation were computed using the corresponding functions from the SEACells package for different latent spaces (Supplementary Fig. 1a-b). Different latent spaces correspond to the diffusion components embedding with different number of dimensions (from 8 to 26).

For the correlation between compactness and separation (Supplementary Fig. 1c), the values computed on 10 diffusion components were used.

### Correlation between metacell size and number of detected genes (Fig. 5b)

The correlation between the metacell size and the number of detected genes was computed for the PBMC dataset by constructing metacells using SuperCell with a graining level of 10. The metacell size corresponds to the number of single cells in a metacell, the number of detected genes corresponds to the number of genes with at least 1 UMI count in a metacell profile.

### Illustrating datasets of different complexity (Fig. 2e)

To illustrate datasets of different complexity (Fig. 2e), 3 single-cell RNA-seq datasets of the same size were used consisting of T cells, PBMCs and lung cells.

T cells dataset was obtained by selecting T cells from the bone marrow datasets downloaded from the SeuratData package^143^. A standard Seurat pipeline^61^ was applied to a randomly subsampled 5’000 cells to obtain a UMAP^95^ representation of data. Cells are colored according to the provided cell type annotation.

The PBMC dataset was expanded with some random noise to 5’000 cells to have the same size as the other two datasets.

The lung dataset was downloaded as a demo Seurat object from (https://seurat.nygenome.org/hlca_ref_files/ts_opt.rds). A standard Seurat pipeline^61^ was applied to a randomly subsampled 5’000 cells to obtain a UMAP^95^ representation of data. Cells are colored according to the provided cell-type annotation.

### Illustrative cartoons (Fig. 1; Fig. 3; Fig. 5a,c; Fig. 6; Fig. 7a-b; Fig.8)

For illustration purposes, no underlying data or real computation were used.

### Code availability

The code to reproduce Figs. 2, 6b, 7c-d, and Supplementary Fig. 1 are available at https://github.com/mariiabilous/Metacell_review_analysis. The code of the tutorial on constructing and analyzing metacells is available at https://github.com/GfellerLab/MetacellAnalysisTutorial. The code of the metacell toolkit for construction and exploration of metacells is available at https://github.com/GfellerLab/MetacellAnalysisToolkit.

## Acknowledgements

We acknowledge the Swiss National Science Foundation Project Grant (31003A_173156). We thank Santiago J. Carmona for providing useful feedback to the manuscript.

