## Supplementary Figure 1 for "Building and analyzing metacells in single-cell genomics data"

### List of supplementary data:

**Supplementary Figure 1. Compactness and separation are latent-space dependent and correlated to each other.**

**Supplementary Table 1: List of studies that introduced methods to build metacells, either as their main aim or as a way to accelerate analyses of large data.**

**Supplementary table 2. Graining levels used across studies.**

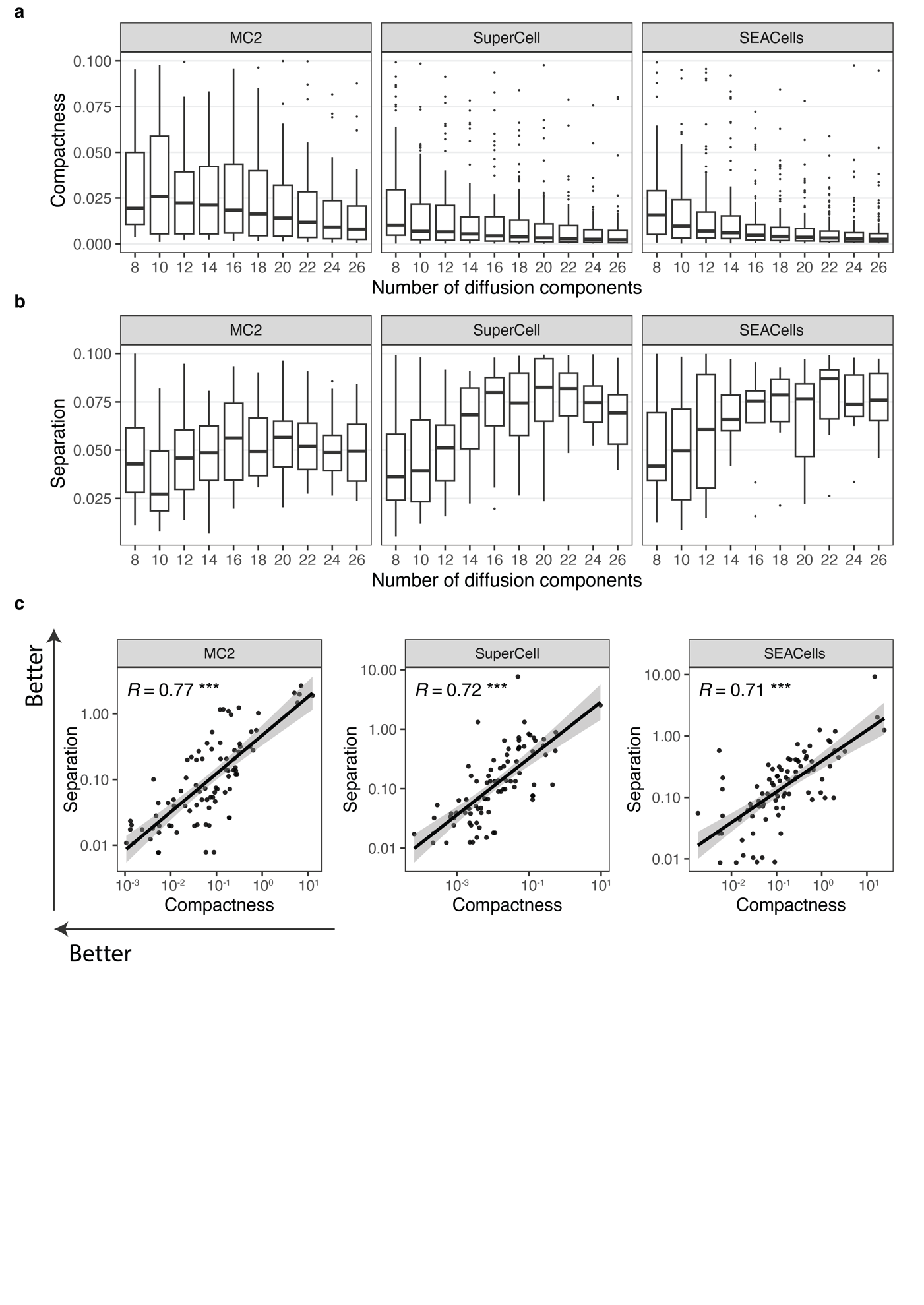

**Supplementary Figure 1. Compactness and separation are latent-space dependent and correlated to each other.**

**a**, Compactness computed at different numbers of diffusion components. **b**, Separation computed at different numbers of diffusion components. **c**, Correlation between metacell compactness and separation. Metacell partition was computed for a PBMC dataset using different metacell construction tools at a graining level of 30. Compactness and separation were computed in the diffusion component space.
